# Systems-level proteomic reprogramming reveals mitochondrial restoration and inhibition of Rho GTPase-mediated cytoskeletal and inflammatory signaling in CKD

**DOI:** 10.64898/2026.09.02.748845

**Authors:** Reshma Shettigar, Shobha Dagamajalu, Hemashree Kampa, Deeksha Salian, Mohd. Altaf Najar

**Affiliations:** Yenepoya Ayurveda Medical College and Hospital,Yenepoya(Deemed to be University), Mangalore 575018, India; Center for Systems Biology and Molecular Medicine, Yenepoya research Center, Yenepoya (Deemed to be University), Mangalore 575018, India

**Keywords:** Chronic kidney disease, Data-independent acquisition proteomics, Dooshivishari Agada, Mitochondrial dysfunction, Oxidative stress, Inflammation

## Abstract

Chronic kidney disease (CKD) is a progressive disorder characterized by metabolic dysfunction, mitochondrial impairment, oxidative stress, and chronic inflammation, ultimately leading to irreversible renal damage. Despite advances in understanding CKD pathophysiology, effective therapies targeting these interconnected molecular processes remain limited. In this study, we performed a comprehensive data-independent acquisition (DIA)-based proteomic analysis to investigate the molecular alterations associated with CKD and to evaluate the therapeutic impact of DVA treatment. Using a CKD model with three treatment conditions (DVA, KY, and DVA+KY) alongside disease and healthy controls, we quantified global proteomic changes and applied statistical filtering (fold change ≥2, p ≤0.05) followed by K-means clustering (k=10). Distinct protein clusters revealed bidirectional modulation upon DVA treatment. Notably, Cluster 1 comprised proteins downregulated in CKD but significantly restored following DVA administration, while Cluster 2 included proteins elevated in CKD that were suppressed by DVA. Pathway enrichment and network analyses demonstrated that Cluster 1 proteins were predominantly associated with mitochondrial function, oxidative phosphorylation, and metabolic processes, whereas Cluster 2 proteins were enriched in immune signaling, oxidative stress, cytoskeletal remodeling, and proteostasis pathways. At the molecular level, DVA treatment restored key mitochondrial and metabolic regulators, including components of the electron transport chain (e.g., COX5A, NDUFS5, SDHB) and redox homeostasis proteins, indicating recovery of cellular bioenergetics. Concurrently, DVA suppressed inflammatory mediators (STAT2, IFI47, GBP2), oxidative stress-related proteins (CYBB, PRDX5), and cytoskeletal regulators linked to renal injury (ARHGEF12, FMNL2). Network and Reactome analyses further confirmed coordinated modulation of interconnected biological systems rather than isolated protein changes.

Collectively, our findings demonstrate that DVA exerts a dual therapeutic effect by restoring essential mitochondrial and metabolic pathways while simultaneously suppressing inflammation, oxidative stress, and cytoskeletal dysregulation in CKD. This systems-level proteomic reprogramming highlights DVA as a promising candidate for CKD intervention and provides mechanistic insights into disease progression and therapeutic targeting.

## Introduction

Chronic kidney disease (CKD) is a global health burden affecting millions of individuals worldwide and is associated with high morbidity and mortality [1]. It is characterized by a progressive decline in renal function, ultimately leading to end-stage renal disease (ESRD) and the need for dialysis or transplantation [2]. The pathogenesis of CKD is complex and multifactorial, involving a combination of metabolic dysregulation, oxidative stress, chronic inflammation, and structural remodeling of renal tissue [3]. Despite advances in clinical management, current therapeutic strategies primarily focus on slowing disease progression rather than reversing underlying molecular damage, highlighting the need for a deeper mechanistic understanding of CKD [4].

Traditional medicine systems have employed herbal medicines, dietary modifications, and lifestyle interventions for different disease conditions. Ayurvedic formulations comprise diverse bioactive compounds, including alkaloids, flavonoids, glycosides, and essential oils. While these therapeutic effects are attributed to synergistic, multi-target pharmacological mechanisms [5], the specific molecular pathways driving their efficacy in chronic kidney disease (CKD) remain poorly characterized [6]. DooshivishariAgada (DVA) is a classical herbomineral formulation traditionally indicated for Dooshivisha (cumulative toxicity), which is responsible for pathological damage mediated by oxidative stress and systemic inflammation. The constituents of DVA exhibit anti-inflammatory, antioxidant, immunomodulatory, and hepatoprotective activities, indicating potential therapeutic relevance for CKD. These polyherbalAyurvedic formulations have demonstrated therapeutic potential against multiple pathological processes implicated in CKD pathogenesis. Their diverse pharmacological activities, including immunomodulatory, antioxidant, and anti-inflammatory effects, may act synergistically to target multiple disease mechanisms. However, their mechanisms of action at the molecular level and the effects of their combination have not been systematically investigated.

A growing body of evidence has identified mitochondrial dysfunction as a central driver of CKD progression [7]. Renal tubular cells are highly energy-dependent and rely heavily on mitochondrial oxidative phosphorylation for ATP production. Disruption of mitochondrial function leads to impaired energy metabolism, increased production of reactive oxygen species (ROS), and activation of pro-fibrotic signaling pathways [8]. Several studies have demonstrated reduced expression of electron transport chain components and compromised mitochondrial integrity in CKD, linking bioenergetic failure to renal injury and fibrosis [9]. In parallel, defects in fatty acid oxidation and central carbon metabolism further exacerbate energy deficiency, contributing to tubular atrophy and disease progression [10].

In addition to metabolic dysfunction, chronic inflammation and immune activation play a critical role in CKD pathophysiology [11]. Persistent activation of innate and adaptive immune pathways, including cytokine signaling and interferon responses, promotes tissue damage and fibrosis [12]. Interferon-stimulated genes and immune regulators such as STAT family proteins have been shown to contribute to sustained inflammatory signaling in renal disease [13]. Furthermore, endotoxin-mediated pathways and complement activation amplify immune responses, creating a self-perpetuating cycle of inflammation and injury [14, 15]. Another emerging hallmark of CKD is oxidative stress, driven by an imbalance between ROS production and antioxidant defense mechanisms. Elevated activity of NADPH oxidases, particularly NOX2, and dysregulation of glutathione metabolism have been implicated in renal oxidative damage [16]. Although antioxidant systems are often upregulated as a compensatory response, persistent oxidative stress leads to protein damage, lipid peroxidation, and activation of apoptotic and fibrotic pathways [17].

Beyond metabolic and inflammatory alterations, cytoskeletal remodeling and Rho GTPasesignaling have gained increasing attention in CKD, particularly in the context of podocyte injury and fibrosis [18]. RhoA-mediated signaling regulates actin cytoskeleton dynamics, cellular contractility, and extracellular matrix deposition. Dysregulation of cytoskeletal proteins and actin-associated regulators contributes to loss of cellular architecture, impaired filtration barrier function, and progression of glomerular disease [19]. In addition, alterations in vesicle trafficking and proteostasis further disrupt cellular homeostasis, exacerbating renal dysfunction [19]. While these individual processes have been studied extensively, CKD is fundamentally a systems-level disease, where multiple pathways interact in a highly coordinated manner [20]. Traditional approaches focusing on single genes or pathways provide limited insight into this complexity [21]. In this context, proteomics offers a powerful platform to capture global molecular changes, particularly as protein abundance and activity are directly linked to cellular function and are influenced by post-transcriptional and post-translational regulation. Data-independent acquisition (DIA)-based proteomics, in particular, enables high-depth, reproducible quantification of the proteome and is well suited for studying complex diseases such as CKD [22]. Despite these advances, a critical gap remains in understanding how therapeutic interventions modulate the CKD proteome at a systems level, particularly whether key pathological signatures such as mitochondrial dysfunction and inflammatory activation can be reversed. Addressing this gap is essential for identifying effective therapeutic strategies and uncovering molecular targets for intervention.

In the present study, we employed DIA-based quantitative proteomics to comprehensively profile protein expression changes in a CKD model and to evaluate the therapeutic effects of DVA, KY(a trial drug), and their combination (DVA+KY). By integrating clustering, pathway enrichment, network analysis, and targeted protein validation, we aimed to define the molecular architecture of CKD and to determine how it is reprogrammed upon treatment. Our findings reveal a coordinated, bidirectional proteomic response in which DVA KY, and their combination differentially modulate metabolic, mitochondrial, inflammatory, and cytoskeletal pathways, with the combination therapy demonstrating enhanced restoration of key disease signatures. DVA treatment restores metabolic and mitochondrial pathways while suppressing inflammation, oxidative stress, and cytoskeletal remodeling. These results provide new mechanistic insights into CKD pathogenesis and highlight key molecular axes that can be targeted for therapeutic intervention.

## Materials and Methods

### Materials

Ethylenediaminetetraacetic acid (EDTA, cat no.E4884-100G), sodium dodecyl sulfate (SDS, Cat no. L3771-1KG), guanidine hydrochloride (GuHCl, Cat no.50940-100G), hydrochloric acid (HCl, Cat no. H1758), acetic acid (Cat no. 1.93002.2521, EMPARTA), formic acid (EMSURE, Cat no. 1.00264), Tris-HCl (Cat no. 648317-100G), and triethylammonium bicarbonate (TEAB, Cat no. T7408-500ML) used for demineralization, protein extraction (Probe Sonicator, LABMAN, L53153), and digestion were obtained from Sigma-Aldrich (St. Louis, MO, United States). Dithiothreitol (DTT, Cat no. D9779-25G), iodoacetamide (IAA, Cat no. I6125-10G), and protease inhibitor cocktail (Cat no: 4693159001) were purchased from Sigma Aldrich(St. Louis, MO, United States). Sequencing-grade modified trypsin (Cat no.V5111)was obtained from Promega (Madison, WI, United States). Pepsin (porcine gastric mucosa, Cat no. V195A) used for collagen-enriched extraction was purchased from Sigma-Aldrich (St. Louis, MO, United States). LC–MS grade water (Cat no. W6-4, 4 L), acetonitrile (Cat no.A955-4, 4L), and formic acid(Cat no. 1.00264) were obtained from Fisher Scientific (Fair Lawn, NJ, United States). Solid phase extraction disk (Cat no. 66883-U,) used in stage tip method for peptide cleanup were obtained fromThermo Fisher Scientific (Waltham, MA, United States). Protein concentration was determined using the Pierce BCA Protein Assay Kit (Cat no. 23225) and peptideconcentration was measured using the Pierce Quantitative Colorimetric Peptide Assay(Cat no. 23275) using standard spectrophotometric or colorimetric assays. Mass spectrometric analysis was performed using an Orbitrap Fusion Tribrid mass spectrometer coupled to a EasyLC1200 ultra-high-performance liquid chromatography (UHPLC) system (Thermo Fisher Scientific, Bremen, Germany).

### Methods

#### Animal model generation and experimental design

All animal experiments were conducted in accordance with the institutional ethical guidelines and were approved by the Institutional Animal Ethics Committee (IAEC) of Yenepoya (Deemed to be University). Male Wistar rats (8–10 weeks old, weighing 250–350 g) were used for the study. Chronic kidney disease (CKD) was induced by administering a diet containing 0.75% adenine for 4 weeks. Animals were randomly allocated into five experimental groups (n = 5 per group), following the Organisation for Economic Co-operation and Development (OECD) guidelines. Group 1 served as the normal control and received a standard laboratory diet and water ad libitum without any treatment. Group 2 consisted of untreated CKD animals. Group 3 included CKD animals treated with DooshivishariAgada (DVA) for 4 weeks. Group 4 received a combination of DooshivishariAgada (DVA) and Kapitta Yoga (KY) aqueous solution for 4 weeks, while Group 5 received Kapitta Yoga (KY) aqueous solution alone for the same duration. The doses of DVA and KY were calculated from the corresponding human therapeutic doses using the standard body surface area conversion method. The animal equivalent dose (AED) was determined using the equation: AED (mg/kg) = Human Equivalent Dose (mg/kg) × 6.17, where 6.17 represents the rat-specific conversion factor. Following CKD induction, the respective treatments were administered orally once daily for 4 weeks. At the end of the treatment period, the animals were euthanized, and kidney tissues were collected for subsequent biochemical, histopathological, and proteomic analyses.

#### Tissue collection and processing for proteomics

Kidney tissues were harvested immediately after euthanasia, washed with ice-cold phosphate-buffered saline (PBS) to remove blood contaminants, and snap-frozen in liquid nitrogen. Samples were stored at −80°C until further processing. For proteomic analysis, tissues were homogenized in lysis buffer containing [e.g., 8 M urea, 50 mMTris-HCl (pH 8.0), 1% SDS] supplemented with protease and phosphatase inhibitors. Homogenization was performed using probe sonicator, followed by centrifugation at 14,000 × g for 15 min at 4°C to remove debris. The supernatant containing soluble proteins was collected for further analysis. Protein concentration was determined using BCA, and equal amounts of protein from each sample were used for downstream processing.

#### Protein digestion and peptide preparation

Protein samples were reduced with [e.g., 5 mMdithiothreitol (DTT)] at 56°C for 30 min and alkylated with [e.g., 15 mMiodoacetamide (IAA)] in the dark at room temperature for 30 min. Samples were then diluted to reduce urea concentration below 2 M. Proteins were digested using sequencing-grade trypsin at a ratio of 1:50 (enzyme:protein) and incubated overnight at 37°C. Following digestion, peptides were acidified using formic acid and desalted using [C18 solid-phase extraction columns]. Eluted peptides were dried using a vacuum concentrator and reconstituted in [0.1% formic acid] prior to mass spectrometry analysis.

#### LC-MS/MS data acquisition (DIA proteomics)

Peptide samples were analyzed using an EASY-nLC 1200 ultra-high-performance nano-liquid chromatography system (Thermo Fisher Scientific, Bremen, Germany) coupled online to an OrbitrapFusion Tribrid mass spectrometer (Thermo Fisher Scientific) equipped with a nano-electrospray ionization source. Peptides were separated on an EASY-Spray PepMap RSLC C18 analytical column (2 μm particle size, 100 Å pore size, 50 μm × 15 cm; Thermo Fisher Scientific) using a binary solvent system consisting of 0.1% formic acid in water (solvent A) and 0.1% formic acid in acetonitrile (solvent B). Peptide separation was achieved using a linear gradient of increasing solvent B over the chromatographic run. The eluting peptides were directly introduced into the mass spectrometer via nano-electrospray ionization.

Mass spectrometric analysis was performed in data-independent acquisition (DIA) mode using predefined isolation windows spanning the peptide mass range. Full MS (MS1) scans and DIA MS/MS (MS2) spectra were acquired at high resolution in the Orbitrap analyzer, enabling comprehensive peptide identification and accurate label-free quantification across all samples.

#### Proteomic data processing and quantification

Raw DIA data were processed using DIA-NN software for peptide identification and quantification. Spectral libraries were generated using library-free mode. Protein identification was performed against the UniProt Human protein database, and false discovery rate (FDR) was controlled at 1% at both peptide and protein levels. Protein intensities were normalized across samples, and only proteins consistently quantified across all samples were included for downstream analysis. Differential expression analysis was performed using a fold-change cutoff of ≥2 and p-value ≤0.05. Proteins meeting these criteria were considered significantly regulated.

#### Statistical and multivariate analysis

Data normalization and statistical analyses were performed using Perseus data was normalized using total peptide amount. Unsupervised hierarchical clustering was carried out to assess sample grouping and reproducibility. Pearson correlation analysis was used to evaluate intra- and inter-group similarity. Principal component analysis (PCA) was performed to visualize global proteomic variation across experimental groups. Self-organizing map (SOM) analysis and dendrogram clustering were used to further explore expression patterns and sample relationships.

#### K-means clustering analysis

To identify coordinated protein expression patterns, K-means clustering was performed on differentially expressed proteins using k = 10 clusters. This approach enabled grouping of proteins with similar expression trends across experimental conditions.

Clusters exhibiting biologically relevant patterns—particularly those showing reversal trends upon DVA treatment were selected for further analysis and interpretation.

#### Functional enrichment and pathway analysis

Gene Ontology (GO), KEGG, and Reactome pathway enrichment analyses were performed using [tools: e.g., DAVID, Metascape, Enrichr, or ReactomePA-specify]. Enriched pathways were identified based on statistical significance (p-value < 0.05). Sankey plots were generated to visualize the relationship between proteins and enriched pathways. Proteins from selected clusters were further analyzed to identify key biological processes associated with CKD and treatment response.

#### Protein-protein interaction (PPI) network analysis

Protein-protein interaction networks were constructed using the STRING database and visualized using Cytoscape. Networks were analyzed to identify highly connected nodes and functional modules. Separate networks were generated for proteins restored upon treatment and those suppressed following treatment, allowing identification of key regulatory hubs associated with disease and recovery.

#### Validation of representative proteins

Selected proteins from key biological pathways were further analyzed for expression patterns across experimental groups. Protein abundance values were extracted from proteomic data and visualized using box plots. Comparisons were made between healthy control (G9), CKD (G8), and DVA-treated (G1) groups to validate trends observed in global proteomic analysis. Statistical significance was assessed using [e.g., Student’s t-test or ANOVA].

## Results

### CKD is associated with profound proteomic reprogramming characterized by metabolic dysfunction and inflammatory activation

Chronic kidney disease (CKD) is a progressive disorder marked by persistent loss of renal function and is strongly associated with metabolic dysregulation, mitochondrial dysfunction, and chronic inflammation. Accumulating evidence suggests that impaired energy metabolism, particularly defects in mitochondrial oxidative phosphorylation and fatty acid oxidation, plays a central role in CKD progression. In parallel, sustained activation of inflammatory and stress-response pathways contributes to tissue injury, fibrosis, and functional decline. Despite these advances, a systems-level understanding of how these processes are coordinated at the proteomic level, and how they can be therapeutically modulated, remains incomplete.

Recent studies have highlighted the importance of proteome-wide analyses in uncovering disease mechanisms that are not fully captured at the transcriptomic level, particularly in complex disorders such as CKD where post-transcriptional and post-translational regulation play critical roles. However, comprehensive quantitative proteomic profiling of CKD in the context of therapeutic intervention remains limited. In particular, the extent to which key pathological signatures—such as mitochondrial dysfunction and inflammatory activation can be reversed at the proteome level has not been systematically explored.

To address this gap, we employed data-independent acquisition (DIA)-based quantitative proteomics to comprehensively profile protein expression across healthy controls, CKD disease models, and treatment groups as detailed in figure 1. This approach enabled high-depth, reproducible quantification of the proteome, allowing us to capture global alterations associated with disease progression and therapeutic response.

**Figure 1.**
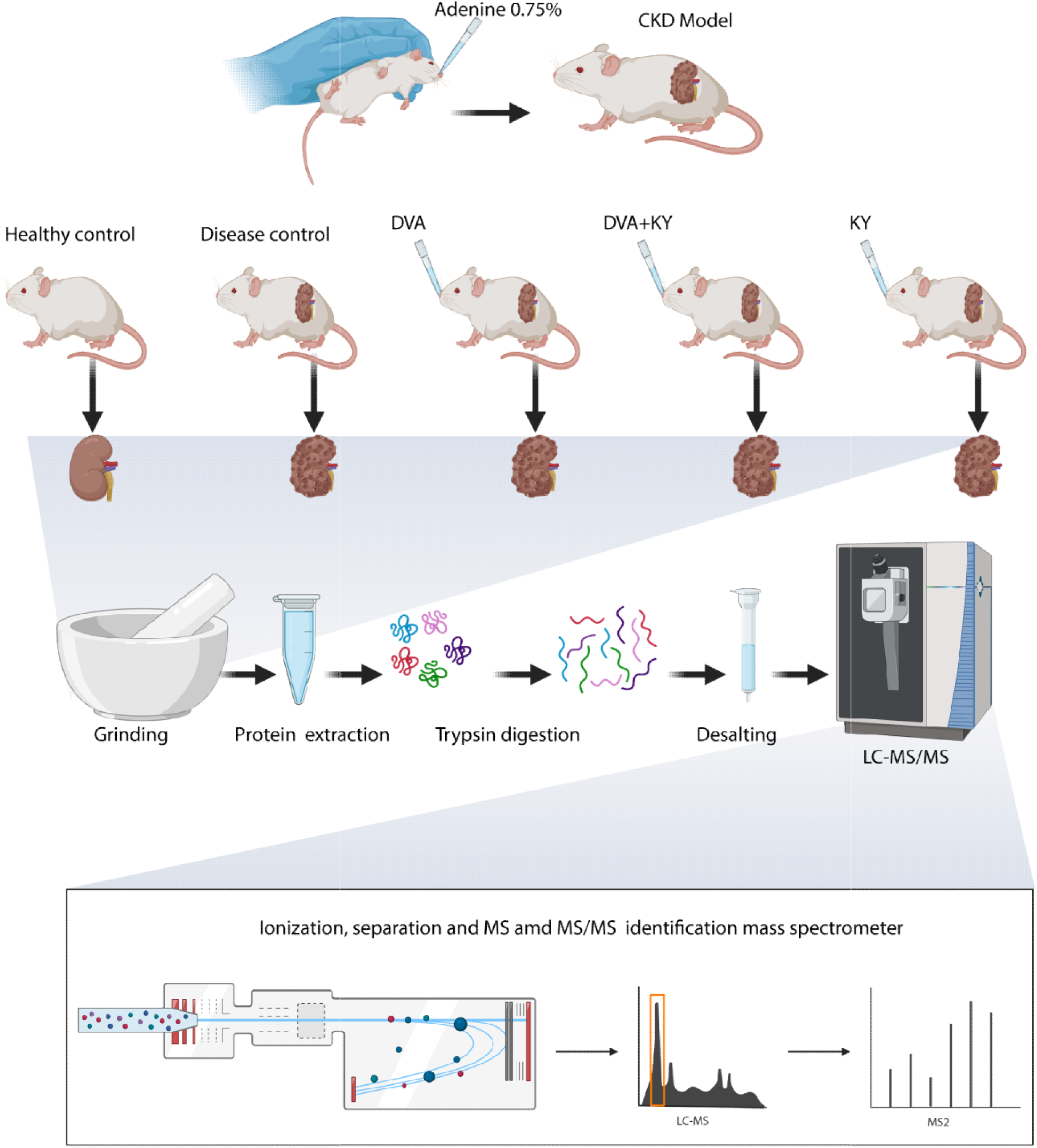
Experimental design and proteomics workflow: Schematic overview of the CKD model, treatment groups (DVA, KY, and combination), and downstream proteomic workflow. Kidney tissues were collected, proteins extracted and digested into peptides, followed by LC-MS/MS analysis using DIA for global proteome profiling and subsequent data analysis.

Unsupervised analyses revealed a clear separation between CKD and healthy control samples, indicating extensive disease-associated proteomic remodeling. Hierarchical clustering demonstrated distinct grouping of CKD samples away from controls, reflecting strong intra-group consistency and pronounced inter-group differences (Figure 2A). This pattern was further supported by correlation analysis, where CKD samples exhibited high internal similarity but reduced correlation with healthy controls (Figure 2B). Principal component analysis (PCA) confirmed this segregation, with CKD samples forming a distinct cluster from controls along the principal components, highlighting global divergence in protein expression profiles (Figure 2C). Similar trends were observed in self-organizing map (SOM) analysis and hierarchical dendrogram clustering, further supporting the robustness of sample classification and underlying proteomic structure (Figure 2D–E). Differential expression analysis identified a substantial number of significantly altered proteins in CKD compared to controls (fold change ≥ 2, p < 0.05), indicating widespread proteome remodeling. Volcano plot visualization revealed a large cohort of both upregulated and downregulated proteins, underscoring the magnitude of disease-associated changes (Supplementary Figure 1A).

**Figure 2.**
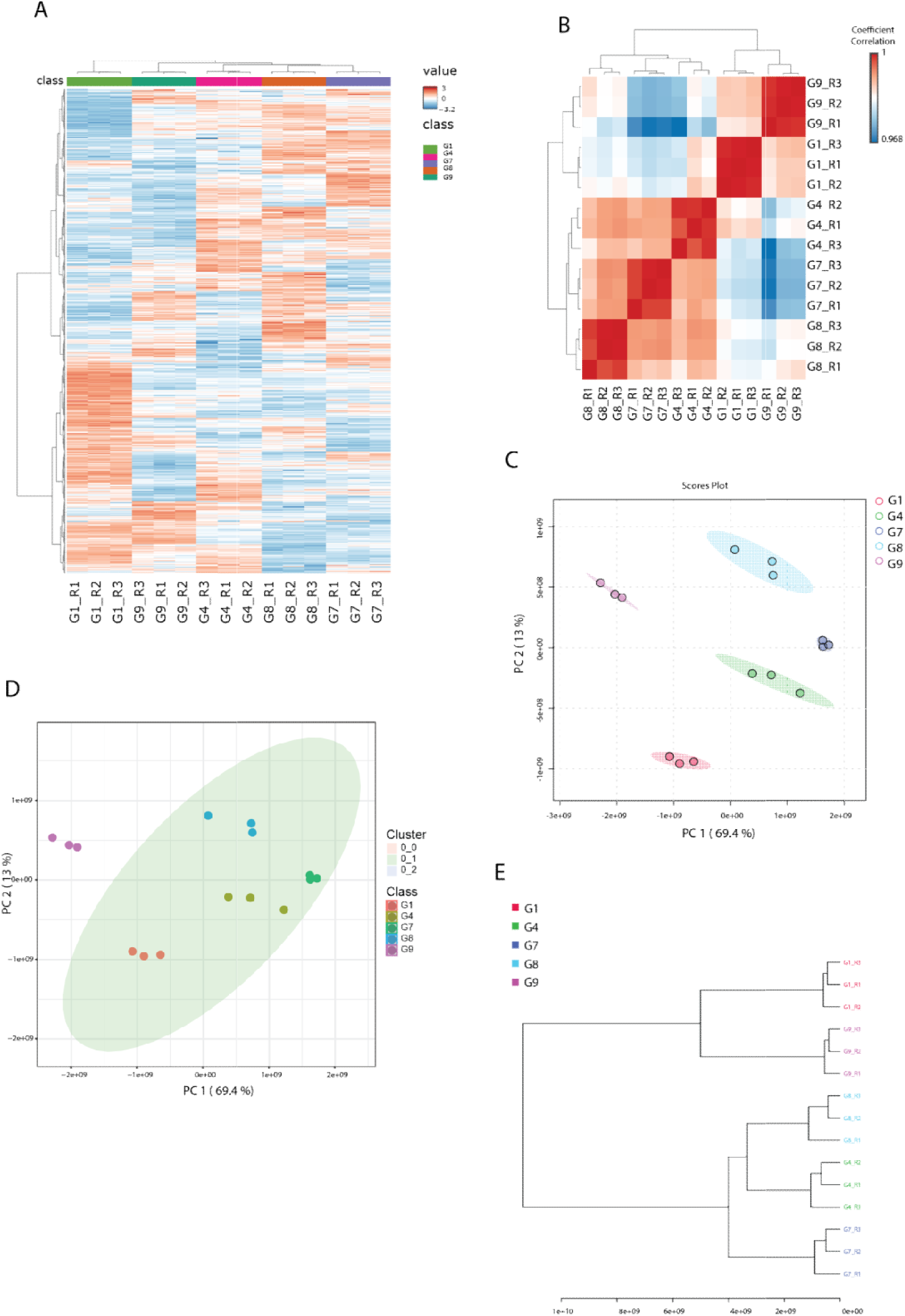
Global proteomic profiling and sample clustering across experimental groups: (A) Heatmap of differentially expressed proteins across all samples, showing distinct expression patterns between healthy control, CKD, and treatment groups. (B) Sample-to-sample correlation heatmap demonstrating high reproducibility within groups and clear separation between conditions. (C) Principal component analysis (PCA) showing clustering of biological replicates and separation of experimental groups. (D) Multivariate clustering plot illustrating distinct grouping of samples based on proteomic profiles. (E) Hierarchical clustering dendrogram confirming segregation of control, CKD, and treatment groups based on global protein expression.

Collectively, these findings demonstrate that CKD is characterized by a coordinated proteomic shift involving suppression of mitochondrial bioenergetics alongside activation of inflammatory and stress-related pathways. This dual signature highlights a systems-level reprogramming of cellular function in CKD and provides a framework for evaluating therapeutic strategies aimed at restoring metabolic balance and reducing inflammation.

### DVA treatment reverses CKD-associated proteomic alterations by restoring metabolic proteins and suppressing disease-associated pathways

To investigate the therapeutic impact of DVA on CKD-associated proteomic alterations, we performed unsupervised K-means clustering (k = 10) to group proteins based on their expression dynamics across treatment conditions. This analysis revealed distinct clusters with treatment-responsive behavior, among which Cluster 1 and Cluster 2 emerged as the most biologically informative (Figure 3A). Cluster 1 comprised proteins that were markedly downregulated in the CKD condition but showed significant restoration following DVA treatment. Visual inspection of the heatmap clearly demonstrated a shift in expression from suppressed levels in CKD toward levels comparable to healthy controls upon DVA administration (Figure 3A). This restoration pattern was further supported by quantitative ratio analysis, where the DVA-treated group (G1) relative to CKD (G8) exhibited a strong increase in protein expression, exceeding even the baseline differences observed between DVA and healthy controls (Figure 3C). This indicates that DVA not only rescues CKD-associated protein loss but may actively enhance recovery toward a healthier proteomic state.

**Figure 3.**
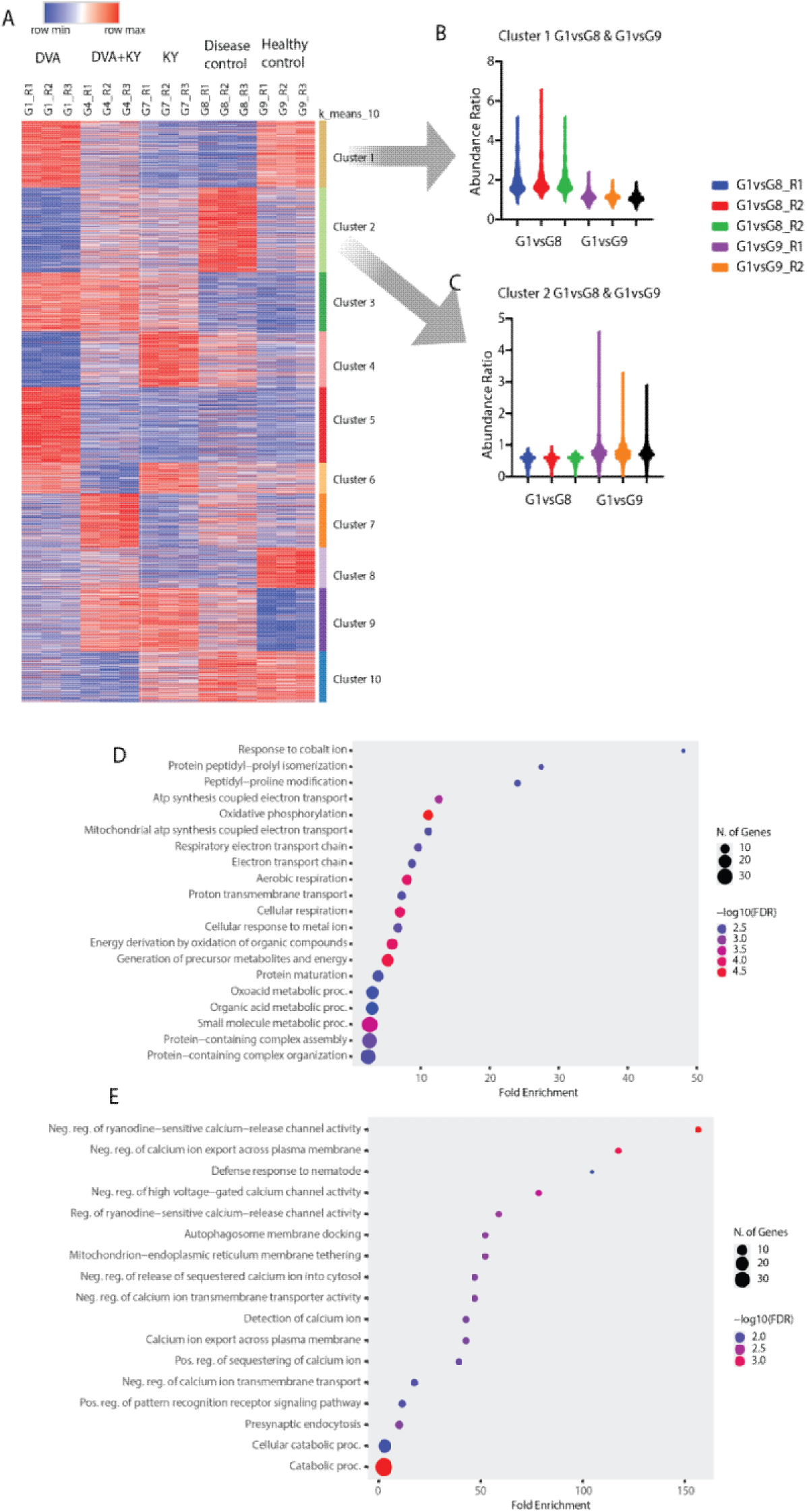
K-means clustering reveals bidirectional proteomic regulation and associated biological processes: (A) Heatmap of differentially expressed proteins grouped by K-means clustering (k = 10), showing distinct expression patterns across DVA, DVA+KY, KY, CKD (disease control), and healthy control groups. (B–C) Representative cluster profiles highlighting Cluster 1 (proteins downregulated in CKD and restored upon DVA treatment) and Cluster 2 (proteins upregulated in CKD and suppressed following DVA treatment). (D) Gene Ontology (GO) biological process enrichment analysis of Cluster 1 proteins, indicating enrichment of mitochondrial function, oxidative phosphorylation, and metabolic processes. (E) GO biological process enrichment analysis of Cluster 2 proteins, showing enrichment of calcium signaling, immune response, vesicle-mediated transport, and cellular stress pathways.

Functional enrichment analysis of Cluster 1 proteins revealed a strong association with mitochondrial and metabolic processes, including oxidative phosphorylation, ATP synthesis, respiratory electron transport, and energy generation pathways (Figure 3D). These findings suggest that DVA treatment effectively restores core bioenergetic functions that are compromised in CKD, highlighting mitochondrial recovery as a central component of its therapeutic action. In contrast, Cluster 2 consisted of proteins that were upregulated in CKD and subsequently suppressed following DVA treatment. The heatmap pattern showed elevated expression in the disease condition, which was markedly reduced upon DVA administration, indicating a reversal of disease-associated protein activation (Figure 3A). Ratio analysis further confirmed this trend, demonstrating that proteins elevated in CKD were significantly downregulated in the DVA-treated group relative to disease controls (Figure 3C).

Biological process enrichment of Cluster 2 proteins revealed pathways associated with CKD pathology, including regulation of calcium signaling, cellular stress responses, autophagy-related processes, and catabolic pathways (Figure 3E). These processes are well known to contribute to renal dysfunction, cellular damage, and disease progression. The observed suppression of these pathways upon DVA treatment suggests a targeted attenuation of pathological signaling networks.

Taken together, these results demonstrate that DVA exerts a dual regulatory effect on the CKD proteome: restoring key metabolic and mitochondrial proteins while simultaneously suppressing disease-associated pathways. This coordinated reprogramming highlights the therapeutic potential of DVA in reversing core molecular features of CKD and underscores the value of cluster-based proteomic analysis in uncovering treatment-responsive biological mechanisms.

### Reactome and network analysis reveal coordinated pathway restoration and suppression following DVA treatment

To further delineate the biological significance of the proteins identified in Cluster 1 and Cluster 2, we performed pathway enrichment analysis using Reactome, followed by network-level interaction mapping. Proteins that were significantly upregulated upon DVA treatment from Cluster 1 and those significantly downregulated from Cluster 2 were independently analyzed to uncover pathway-level reprogramming.

Reactome pathway mapping visualized through Sankey plots revealed a clear segregation of functional processes modulated by DVA treatment (Figure 4A–B). Proteins restored in Cluster 1 predominantly mapped to pathways associated with mitochondrial function and cellular metabolism, including amino acid transport, respiratory electron transport, ATP synthesis, and metabolic processes. These pathways formed a highly interconnected network, indicating a coordinated restoration of bioenergetic and metabolic homeostasis disrupted in CKD (Figure 4A).

**Figure 4.**
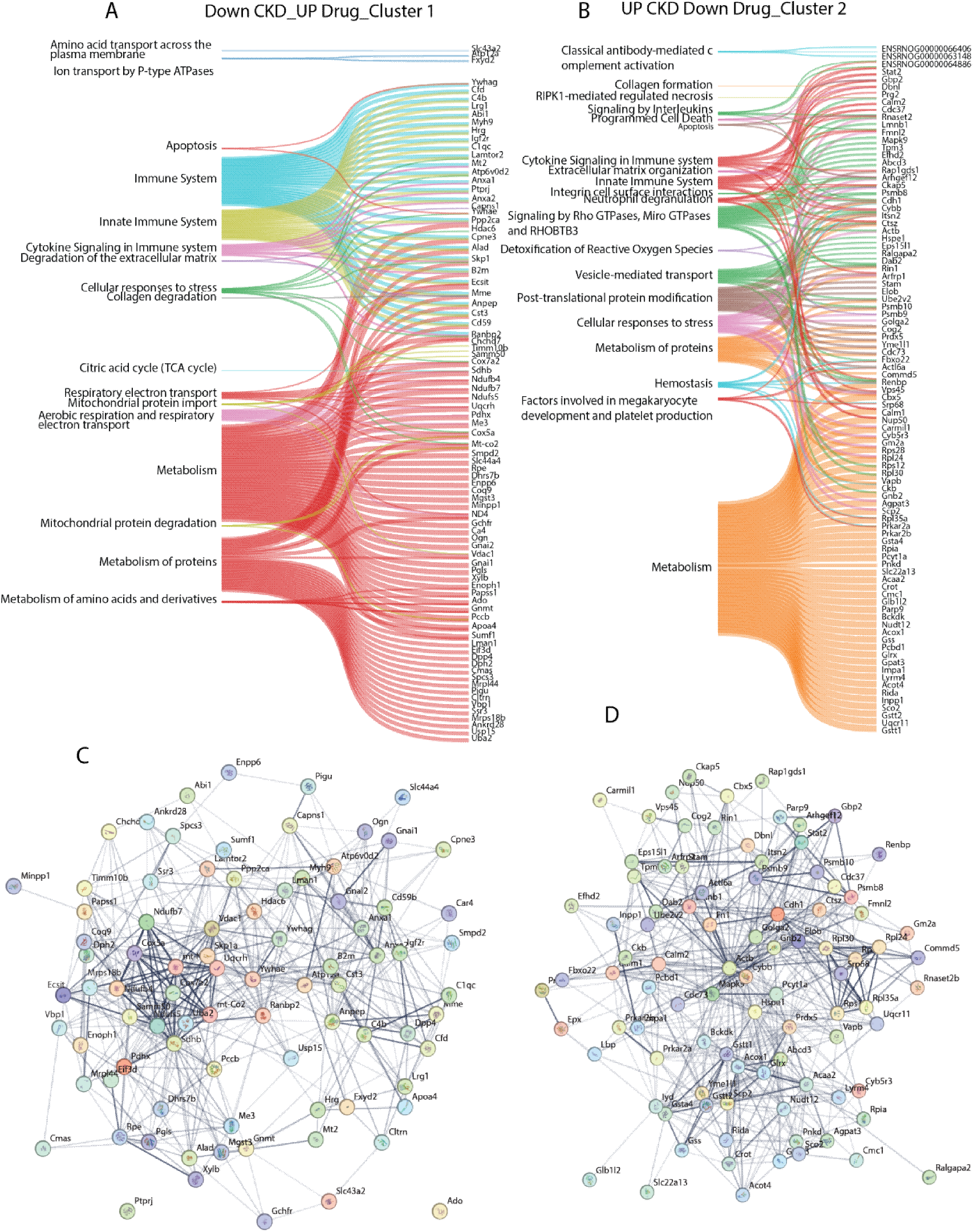
Reactome pathway enrichment and protein–protein interaction networks of DVA-regulated proteins: (A) Sankey plot representing Reactome pathway enrichment of proteins significantly upregulated following DVA treatment (Cluster 1), highlighting pathways associated with metabolism, mitochondrial function, and cellular respiration.(B) Sankey plot showing Reactome pathway enrichment of proteins downregulated following DVA treatment (Cluster 2), indicating suppression of immune signaling, cytokine responses, vesicle-mediated transport, and cellular stress pathways.(C) Protein– protein interaction (PPI) network of upregulated proteins (Cluster 1), illustrating highly interconnected modules associated with metabolic and mitochondrial processes.(D) PPI network of downregulated proteins (Cluster 2), revealing interaction networks linked to immune regulation, cytoskeletal organization, and proteostasis.

In contrast, proteins downregulated in Cluster 2 were enriched in pathways linked to immune activation, extracellular matrix organization, cytokine signaling, and stress-related processes (Figure 4B). Notably, pathways such as complement activation, interleukin signaling, neutrophil degranulation, and collagen formation were prominently represented, highlighting suppression of inflammatory and fibrotic signaling cascades that are central to CKD progression.

To further understand the functional relationships among these proteins, protein–protein interaction (PPI) networks were constructed. The Cluster 1-derived proteins formed a dense interaction network centered around mitochondrial and metabolic hubs, supporting their role in restoring core cellular functions (Figure 4C). Key nodes within this network were associated with oxidative phosphorylation and energy metabolism, reinforcing the concept of mitochondrial recovery as a primary therapeutic effect of DVA.

Conversely, the Cluster 2-derived proteins also exhibited strong interconnectivity, forming networks enriched for immune and extracellular signaling components (Figure 4D). The suppression of these tightly linked protein modules suggests that DVA effectively disrupts coordinated pathological signaling networks rather than acting on isolated targets.

Supporting these findings, extended functional annotations provided in Supplementary Figure 4 further validated the pathway enrichment patterns. Specifically, Supplementary Figure 4A–C corroborated the enrichment of mitochondrial and metabolic processes among upregulated proteins, while Supplementary Figure 4D–F confirmed the downregulation of pathways related to immune response, protein catabolism, and stress signaling.

Collectively, these results demonstrate that DVA induces a systems-level reprogramming of the CKD proteome, characterized by restoration of metabolic and mitochondrial pathways alongside suppression of inflammation- and fibrosis-associated networks. This coordinated modulation of interconnected biological processes provides mechanistic insight into the therapeutic efficacy of DVA in CKD.

### DVA treatment restores mitochondrial, metabolic, and structural protein networks disrupted in CKD

To validate the functional significance of the proteomic alterations identified through clustering and pathway analyses, we examined the expression profiles of selected key proteins representing major biological axes affected in CKD. These included proteins involved in mitochondrial respiration, redox metabolism, epigenetic regulation, cytoskeletal organization, and membrane transport. Comparative analysis across healthy control (G9), CKD (G8), and DVA-treated (G1) groups revealed a consistent pattern in which these proteins were significantly downregulated in CKD and subsequently restored upon DVA treatment (Figure 5).

**Figure 5.**
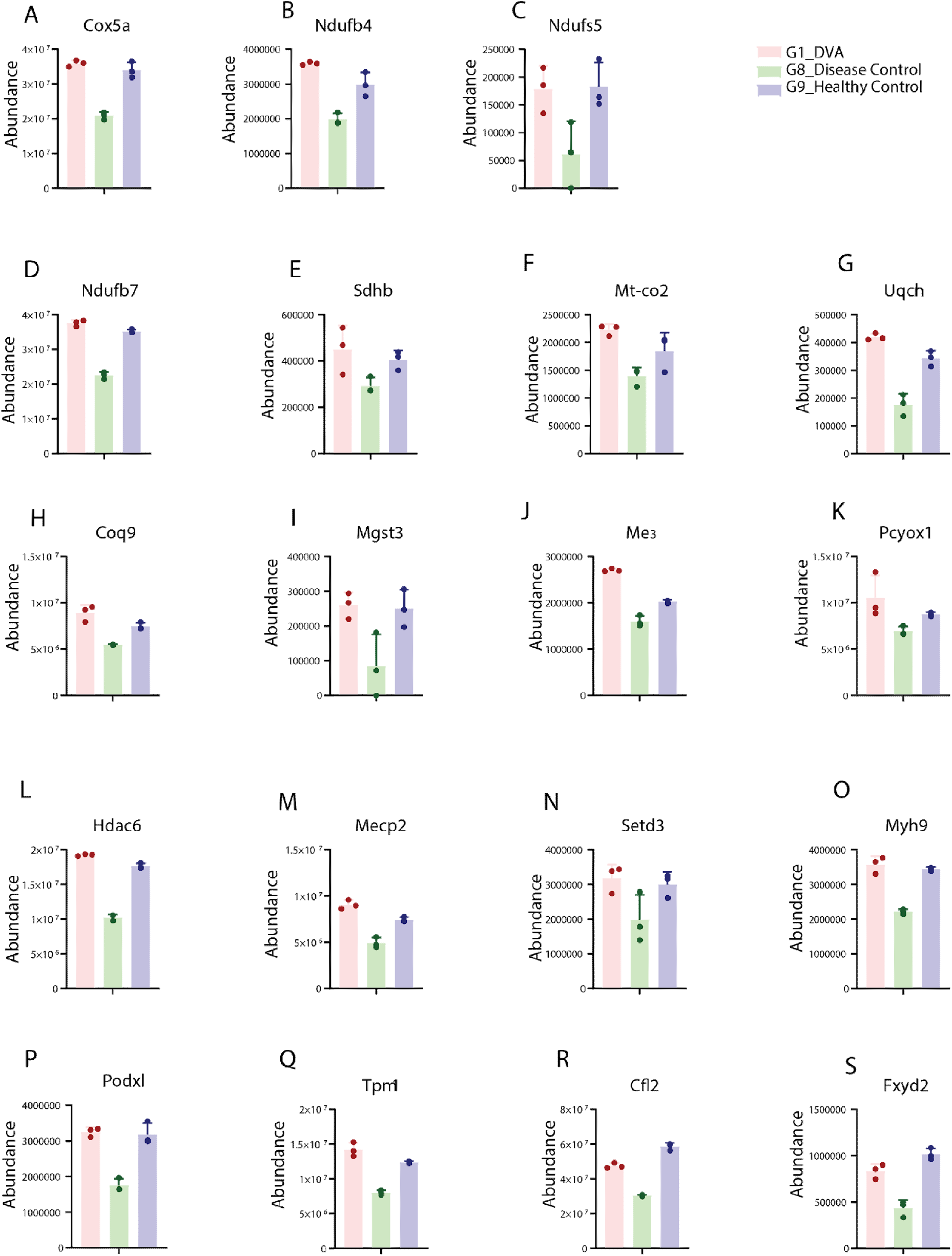
Validation of cluster 2 proteins showing DVA-mediated suppression of CKD-associated pathways: Bar plots depicting the relative protein expression levels of selected cluster 2 proteins across experimental groups: healthy control (G9), CKD (G8), and DVA-treated (G1). Proteins shown are significantly upregulated in CKD and are consistently reduced following DVA treatment, indicating reversal of disease-associated molecular signatures. Data are presented as mean ± SEM with individual data points overlaid

Mitochondrial dysfunction is a central hallmark of CKD, characterized by impaired oxidative phosphorylation and reduced ATP production. Several subunits of the electron transport chain, including Cox5a, Mt-co2, Uqcrh, and Coq9, play essential roles in maintaining mitochondrial respiratory efficiency. Cox5a and Mt-co2 are integral components of cytochrome c oxidase (Complex IV), critical for terminal electron transfer and ATP synthesis, while Uqcrh is involved in Complex III stability and electron flux. Coq9 participates in coenzyme Q biosynthesis, which is indispensable for electron transport chain function. In parallel, Ndufs5, Ndufb4, and Ndufb7, key subunits of Complex I, along with Sdhb, a core component of Complex II, are essential for maintaining mitochondrial electron transport integrity. Consistent with previous reports linking CKD to mitochondrial impairment, these proteins were markedly reduced in the CKD group. Notably, DVA treatment significantly restored their expression levels, indicating recovery of mitochondrial bioenergetic capacity (Figure 5).

Redox homeostasis and metabolic balance are also severely disrupted in CKD. Proteins such as Mgst3 and Gss are critical regulators of glutathione metabolism and cellular antioxidant defense, while Me3 contributes to NADPH generation, supporting redox balance. Pcyox1, involved in oxidative metabolic processes, has been associated with lipid metabolism and oxidative stress regulation. The observed downregulation of these proteins in CKD reflects increased oxidative stress and impaired detoxification capacity, both well-documented features of renal pathology. Restoration of their expression following DVA treatment suggests an enhanced antioxidant response and improved metabolic resilience (Figure 5).

Epigenetic and transcriptional regulators further contribute to CKD progression by modulating gene expression programs associated with fibrosis, inflammation, and cellular stress. Hdac6, a histone deacetylase, has been implicated in cytoskeletal remodeling and stress response pathways, while Mecp2 functions as a transcriptional regulator linking DNA methylation to gene repression. Setd3, a histidine methyltransferase, plays roles in actin dynamics and cellular differentiation. Dysregulation of these proteins in CKD may contribute to maladaptive transcriptional reprogramming. Their normalization upon DVA treatment indicates a potential role for epigenetic modulation in mediating therapeutic effects (Figure 5).

Structural integrity and cellular architecture are also compromised in CKD, particularly in specialized renal cells such as podocytes. Proteins including Myh9, Tpm1, and Cfl2 are central to actin cytoskeleton organization and contractile function, while Podxl (podocalyxin) is a key determinant of podocyte structure and filtration barrier integrity. Loss or dysfunction of these proteins has been directly linked to glomerular injury and proteinuria. In our dataset, these proteins were consistently reduced in CKD and significantly restored following DVA treatment, suggesting recovery of cytoskeletal stability and renal structural integrity (Figure 5).

Finally, membrane transport and ion homeostasis, critical for kidney function, were represented by Fxyd2, a regulatory subunit of Na⁺/K⁺-ATPase. Fxyd2 is essential for maintaining electrolyte balance and tubular transport function, and its dysregulation has been associated with renal dysfunction. The observed decrease in CKD and subsequent restoration with DVA treatment further supports the functional recovery of renal transport mechanisms (Figure 5).

Collectively, these findings demonstrate that DVA treatment restores multiple interconnected protein networks disrupted in CKD, including mitochondrial respiration, redox balance, epigenetic regulation, cytoskeletal organization, and membrane transport. The coordinated recovery of these critical biological systems provides strong mechanistic evidence supporting the therapeutic potential of DVA in reversing CKD-associated molecular dysfunction.

### DVA treatment suppresses CKD-associated inflammatory, oxidative, and cytoskeletal signaling networks

To further define the disease-driving proteomic signatures of CKD and their reversal upon treatment, we analyzed a panel of key proteins derived from Cluster 2 that were significantly upregulated in CKD and subsequently reduced following DVA administration. Comparative expression analysis across healthy control (G9), CKD (G8), and DVA-treated (G1) groups revealed a consistent and robust pattern: these proteins were markedly elevated in CKD and significantly suppressed following DVA treatment, often returning toward baseline levels observed in healthy controls (Figure 6). This confirms that DVA actively counteracts core pathological pathways associated with CKD.

**Figure 6.**
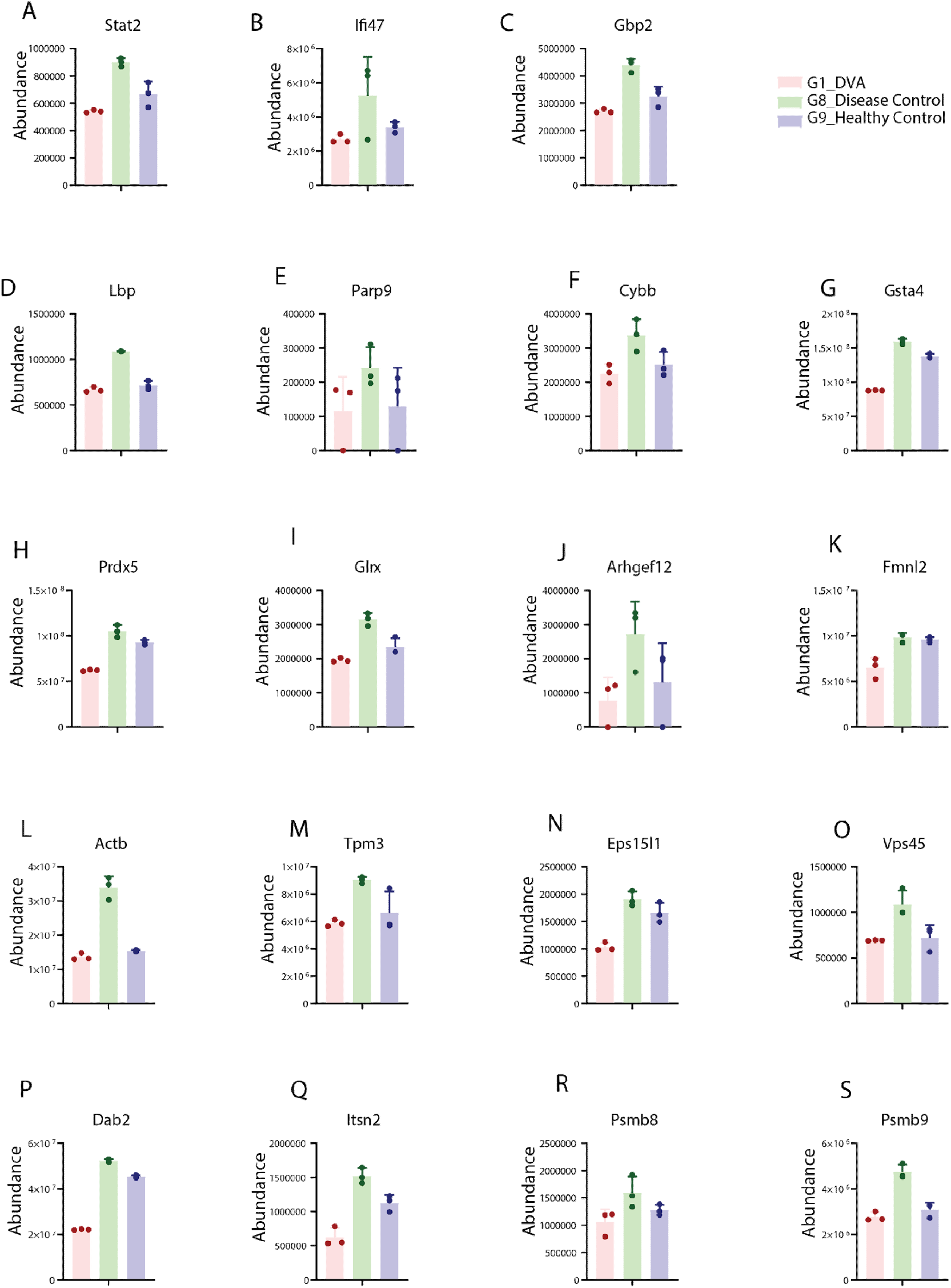
Validation of cluster 1 proteins showing DVA-mediated restoration of mitochondrial and metabolic pathways: Bar plots showing relative expression levels of selected cluster 1 proteins across healthy control (G9), CKD (G8), and DVA-treated (G1) groups. These proteins are significantly downregulated in CKD and are restored toward control levels following DVA treatment. Data are presented as mean ± SEM with individual data points indicated.

A major subset of these proteins is linked to immune and inflammatory signaling, a central hallmark of CKD progression. Stat2, a key transcription factor in type I interferon signaling, is known to amplify inflammatory cascades and contribute to sustained immune activation in renal disease. Similarly, Ifi47 and Gbp2, both interferon-inducible GTPases, are involved in innate immune responses and have been associated with chronic inflammatory states. Lbp, a mediator of endotoxin recognition, plays a critical role in triggering systemic inflammatory signaling, while Parp9 is implicated in interferon-mediated transcriptional responses and immune stress pathways. In agreement with these established roles, our data show a pronounced increase in these proteins in CKD, reflecting heightened inflammatory activation. Importantly, DVA treatment significantly reduced their expression (Figure 6), indicating effective suppression of interferon-driven and innate immune pathways.

Oxidative stress-related proteins were also prominently elevated in CKD and normalized following treatment. Cybb (NOX2) is a major enzymatic source of reactive oxygen species and a well-recognized contributor to renal oxidative injury. Antioxidant and redox-regulating proteins such as Gsta4, Prdx5, and Glrx are typically induced under oxidative stress conditions as compensatory mechanisms. Their increased expression in CKD reflects persistent redox imbalance and cellular stress. The marked reduction of these proteins following DVA treatment (Figure 6) suggests a decrease in oxidative burden and restoration of redox homeostasis.

A particularly important finding from this analysis is the involvement of proteins associated with Rho GTPasesignaling and cytoskeletal remodeling, which represent a critical mechanistic axis in kidney injury. Arhgef12, a Rho guanine nucleotide exchange factor, activates RhoAsignaling and promotes cytoskeletal reorganization, contributing to fibrosis and cellular contractility. Fmnl2, a formin protein, regulates actin filament assembly, while Actb and Tpm3 are fundamental components of the actin cytoskeleton. Dysregulation of these proteins is closely linked to altered cell morphology, increased motility, and fibrotic remodeling in renal cells. In our dataset, these proteins were significantly upregulated in CKD and consistently suppressed following DVA treatment (Figure 6), suggesting that DVA modulates Rho GTPase-driven cytoskeletal signaling and may limit structural and fibrotic damage.

In addition, proteins involved in vesicle trafficking and endocytosis, including Eps15l1, Vps45, Dab2, and Itsn2, were elevated in CKD and reduced upon treatment. These proteins regulate membrane dynamics, receptor internalization, and intracellular transport, processes that are often dysregulated under pathological conditions. Their normalization following DVA treatment indicates restoration of cellular trafficking and signaling balance (Figure 6). Finally, components of the immunoproteasome, including Psmb8 and Psmb9, were significantly upregulated in CKD. These proteins are induced under inflammatory conditions and play key roles in antigen processing and immune activation. Their elevated levels are consistent with chronic inflammatory signaling in CKD. Notably, DVA treatment reduced their expression (Figure 6), further supporting its role in attenuating immune activation and restoring proteostatic balance.

Collectively, these findings demonstrate that DVA treatment exerts a broad suppressive effect on CKD-associated pathological networks, including inflammation, oxidative stress, cytoskeletal remodeling, vesicle trafficking, and proteasome activation. The coordinated downregulation of these interconnected pathways highlights the ability of DVA to counteract key molecular drivers of CKD progression, complementing its restorative effects on mitochondrial and metabolic functions described earlier.

DVA treatment induces a robust and coordinated reversal of CKD-associated proteomic alterations, characterized by simultaneous restoration of mitochondrial and metabolic functions alongside suppression of inflammatory, oxidative, and cytoskeletal remodeling pathways. Proteins involved in oxidative phosphorylation, redox homeostasis, and structural integrity that were significantly reduced in CKD were consistently restored upon DVA treatment, while key mediators of immune activation, ROS generation, Rho GTPasesignaling, and proteostasis that were elevated in disease were markedly suppressed. This bidirectional regulation highlights a systems-level therapeutic effect, wherein DVA not only rescues essential cellular functions but also attenuates core drivers of CKD pathology, underscoring its potential as an effective intervention for restoring renal molecular homeostasis.

## Discussion

Chronic kidney disease (CKD) is a multifactorial disorder characterized by progressive loss of renal function driven by interconnected processes including mitochondrial dysfunction, chronic inflammation, oxidative stress, metabolic dysregulation, and cytoskeletal remodeling[23, 24]. Despite extensive investigation, the molecular mechanisms underlying CKD progression remain incompletely understood, and effective therapeutic strategies targeting these interconnected pathways are limited. In this study, we employed a systems-level proteomic approach to characterize CKD-associated molecular alterations and to evaluate the therapeutic effects of DVA treatment. Our findings reveal that DVA induces a coordinated and bidirectional reprogramming of the CKD proteome, simultaneously restoring essential cellular functions while suppressing disease-promoting pathways.

One of the most prominent features of CKD identified in our study is the widespread suppression of mitochondrial proteins and bioenergetic pathways. Mitochondrial dysfunction is increasingly recognized as a central driver of CKD progression, contributing to impaired ATP production, increased reactive oxygen species (ROS) generation, and tubular injury [25–27]. Previous studies have demonstrated that defects in oxidative phosphorylation and electron transport chain (ETC) components are closely associated with renal fibrosis and decline in kidney function [9, 28]. In line with these observations, we found that multiple ETC components, including subunits of Complex I (NDUFB4, NDUFB7, NDUFS5), Complex II (SDHB), Complex III (UQCRH), and Complex IV (COX5A, MT-CO2), were significantly downregulated in CKD. Restoration of these proteins following DVA treatment suggests recovery of mitochondrial integrity and improved cellular energy metabolism. Additionally, the upregulation of COQ9, involved in coenzyme Q biosynthesis, further supports enhanced electron transport efficiency. Collectively, these findings position mitochondrial restoration as a central therapeutic axis of DVA action. CKD is also characterized by profound metabolic and redox imbalance [29]. Increased oxidative stress, driven by excessive ROS production and impaired antioxidant defenses, plays a critical role in renal injury and progression to fibrosis [30–32]. Proteins such as CYBB (NOX2), a major source of ROS, are known to be elevated in CKD and contribute to oxidative damage [33, 34]. In parallel, antioxidant systems including glutathione metabolism are often dysregulated[35]. Our data demonstrate that proteins involved in redox regulation, including GSTA4, PRDX5, and GLRX, are elevated in CKD, reflecting a compensatory but insufficient response to oxidative stress. DVA treatment significantly reduced the expression of these stress-associated proteins while restoring key antioxidant regulators such as MGST3 and GSS. This suggests that DVA not only reduces oxidative burden but also re-establishes redox homeostasis, a critical factor in preventing renal damage.

Inflammation is another hallmark of CKD, with persistent activation of innate and adaptive immune pathways contributing to disease progression [36]. Interferon signaling, in particular, has been implicated in sustaining chronic inflammation in renal disease [37, 38]. Proteins such as STAT2, IFI47, and GBP2 are key mediators of interferon responses and are associated with immune activation and tissue injury [39]. Additionally, LBP-mediated endotoxin signaling and PARP9-driven immune responses further amplify inflammatory cascades [40]. Our findings demonstrate a marked upregulation of these proteins in CKD, consistent with previous reports. Importantly, DVA treatment significantly suppressed their expression, indicating attenuation of interferon-driven and innate immune signaling. This highlights the anti-inflammatory potential of DVA and its ability to disrupt chronic immune activation in CKD. Beyond metabolic and inflammatory alterations, cytoskeletal remodeling and Rho GTPase signaling have emerged as critical contributors to renal pathology, particularly in podocyte injury and fibrosis. Activation of RhoAsignaling pathways promotes actin cytoskeleton reorganization, increased cellular contractility, and extracellular matrix deposition [41]. ARHGEF12, a key activator of RhoA, along with actin-associated proteins such as FMNL2, ACTB, and TPM3, play essential roles in these processes [42]. Dysregulation of these proteins has been linked to podocyte dysfunction, loss of filtration barrier integrity, and progression of glomerular disease [43, 44]. In our study, these cytoskeletal regulators were significantly upregulated in CKD and suppressed following DVA treatment, suggesting that DVA modulates Rho GTPase-driven cytoskeletal dynamics. Concurrently, restoration of structural proteins such as MYH9, TPM1, CFL2, and PODXL further supports recovery of cellular architecture and renal integrity.

Vesicle trafficking and proteostasis pathways were also significantly altered in CKD. Proteins such as EPS15L1, VPS45, DAB2, and ITSN2 regulate endocytosis and intracellular transport, processes that are often disrupted in disease states and contribute to abnormal signaling and protein accumulation [45]. Similarly, components of the immunoproteasome, including PSMB8 and PSMB9, are upregulated under inflammatory conditions and play roles in antigen processing and immune activation [46]. The suppression of these proteins following DVA treatment suggests restoration of cellular homeostasis and reduced proteotoxic stress. These findings align with emerging evidence that dysregulated proteostasis contributes to CKD progression and that targeting these pathways may offer therapeutic benefit. Importantly, our integrative analysis demonstrates that these molecular alterations are not isolated but occur within highly interconnected networks. Reactome pathway and protein–protein interaction analyses revealed coordinated modulation of entire biological systems, rather than individual proteins. This systems-level reprogramming underscores the multifaceted nature of CKD and highlights the importance of targeting multiple pathways simultaneously. DVA appears to achieve this by acting on central regulatory nodes, particularly mitochondrial function and cytoskeletal signaling, thereby influencing downstream metabolic, inflammatory, and structural processes.

The dual effect of DVA restoring essential cellular functions while suppressing pathological pathways represents a significant therapeutic advantage. Most current CKD treatments primarily focus on slowing disease progression rather than reversing underlying molecular damage. In contrast, our findings suggest that DVA has the potential to actively reprogram the diseased proteome toward a healthier state. This bidirectional mechanism may be critical for achieving meaningful functional recovery in CKD. Despite these promising findings, certain limitations should be acknowledged. The study is based on a preclinical model with a limited sample size, and further validation in larger cohorts and clinical samples is to confirm translational relevance. Additionally, while proteomic analysis provides valuable insights into molecular changes, functional studies are needed to establish causal relationships and to delineate the precise mechanisms of DVA action.

In conclusion, our study provides a comprehensive proteomic framework for understanding CKD pathogenesis and therapeutic response. By integrating clustering, pathway enrichment, network analysis, and targeted protein validation, we demonstrate that DVA induces a coordinated restoration of mitochondrial and metabolic functions while suppressing inflammation, oxidative stress, and cytoskeletal remodeling. These findings not only highlight the therapeutic potential of DVA but also identify key molecular axes that may serve as targets for future CKD interventions.

## Supporting information

Volcano plot visualization revealed a large cohort of both upregulated and downregulated proteins, underscoring the magnitude of disease-associated ch

## Ethics approval and consent to participate

The Institutional Animal Ethics Committee approved the study with a reference number of YU/IAEC/P15/2023.

## Data Availability Statement

Data related to the study are provided in the manuscript and supplementary rest of the data and can be provided request.

## Author contributions

**Mohd. Altaf Najar*:** Conceptualization, proteomic data curation, proteomic data analysis, statistical analysis, and original draft preparation.**Reshma Shettigar*:** Conceptualization, funding acquisition, project administration, supervision, methodology, and final manuscript editing. **Shobha D:** Conceptualization, methodology optimization, and proteomic data acquisition.**Hemashree:** Animal care, dissection, tissue collection, and investigation support.**DeekshaSalian:** Animal care, dissection, tissue collection, and investigation support.

## Abbreviations

CKD: Chronic kidney disease
ESRD: End-stage renal disease
DVA: Dooshivishari Agada
KY: Kapitta Yoga
DIA: Data-independent acquisition
LC-MS/MS: Liquid chromatography–tandem mass spectrometry
UHPLC: Ultra-high-performance liquid chromatography
PBS: Phosphate-buffered saline
BCA: Bicinchoninic acid
DTT: Dithiothreitol
IAA: Iodoacetamide
TEAB: Triethylammonium bicarbonate
EDTA: Ethylenediaminetetraacetic acid
SDS: Sodium dodecyl sulfate
GuHCl: Guanidine hydrochloride
HCl: Hydrochloric acid
FDR: False discovery rate
PCA: Principal component analysis
SOM: Self-organizing map
PPI: Protein–protein interaction
GO: Gene Ontology
KEGG: Kyoto Encyclopedia of Genes and Genomes
ROS: Reactive oxygen species
ETC: Electron transport chain
ATP: Adenosine triphosphate
NADPH: Nicotinamide adenine dinucleotide phosphate (reduced form)
NOX2: NADPH oxidase 2
MAF: Mutation Annotation Format
CNA: Copy-number alteration
GTP: Guanosine triphosphate
RhoA: Ras homolog family member A
OECD: Organisation for Economic Co-operation and Development
IAEC: Institutional Animal Ethics Committee
AED: Animal equivalent dose
SEM: Standard error of the mean

