## Supplementary material for "Systems-level proteomic reprogramming reveals mitochondrial restoration and inhibition of Rho GTPase-mediated cytoskeletal and inflammatory signaling in CKD": Volcano plot visualization revealed a large cohort of both upregulated and downregulated proteins, underscoring the magnitude of disease-associated ch

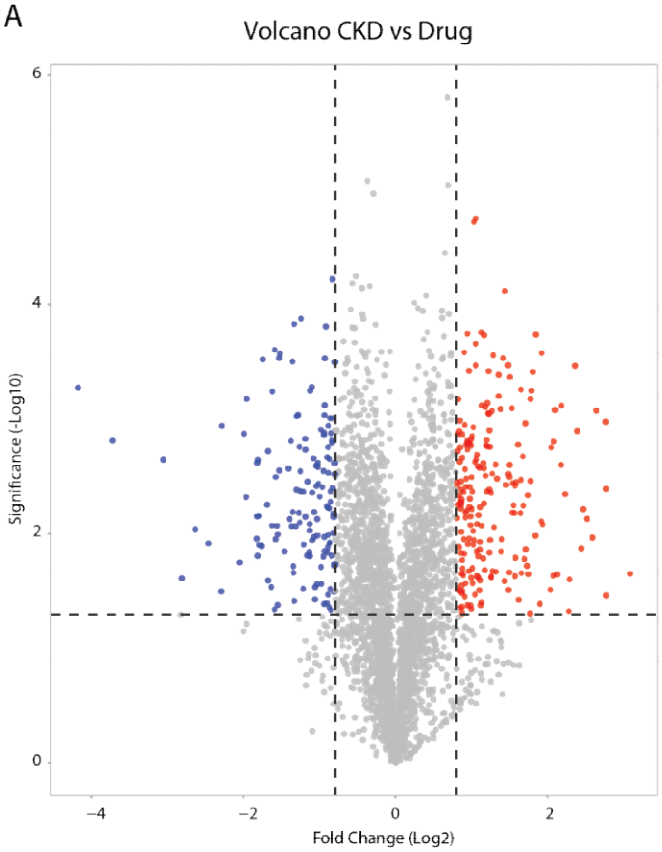
Volcano CKD vs Drug
Significance (-Log10)
Fold Change (Log2)
6
4
2
0
−4
−2
0
2

B

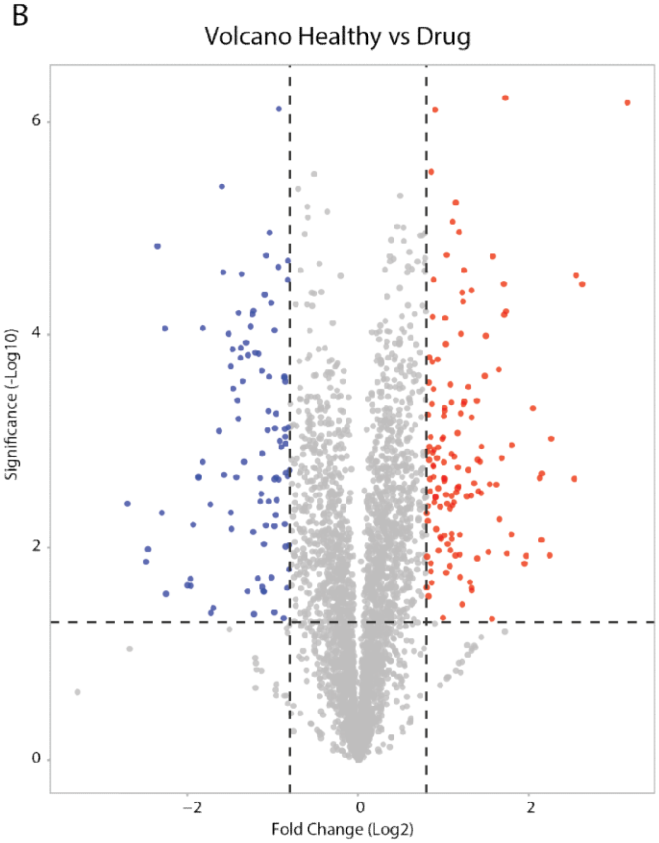
Volcano Healthy vs Drug
Significance (-Log10)
Fold Change (Log2)
6
4
2
0
−2
0
2

**Figure Supplementary 1. Differential proteomic landscape of CKD and DVA treatment:** Volcano plots depicting differentially expressed proteins between experimental groups. The upper panel represents the comparison between CKD (G8) and healthy control (G9), while the lower panel represents CKD (G8) versus DVA-treated (G1) samples. Each dot corresponds to an individual protein, with red indicating significantly upregulated proteins, blue indicating significantly downregulated proteins, and gray representing non-significant changes. Vertical dashed lines indicate fold-change thresholds, and the horizontal dashed line represents the statistical significance cutoff. These plots highlight the widespread proteomic alterations in CKD and the modulatory effect of DVA treatment on disease-associated protein expression.

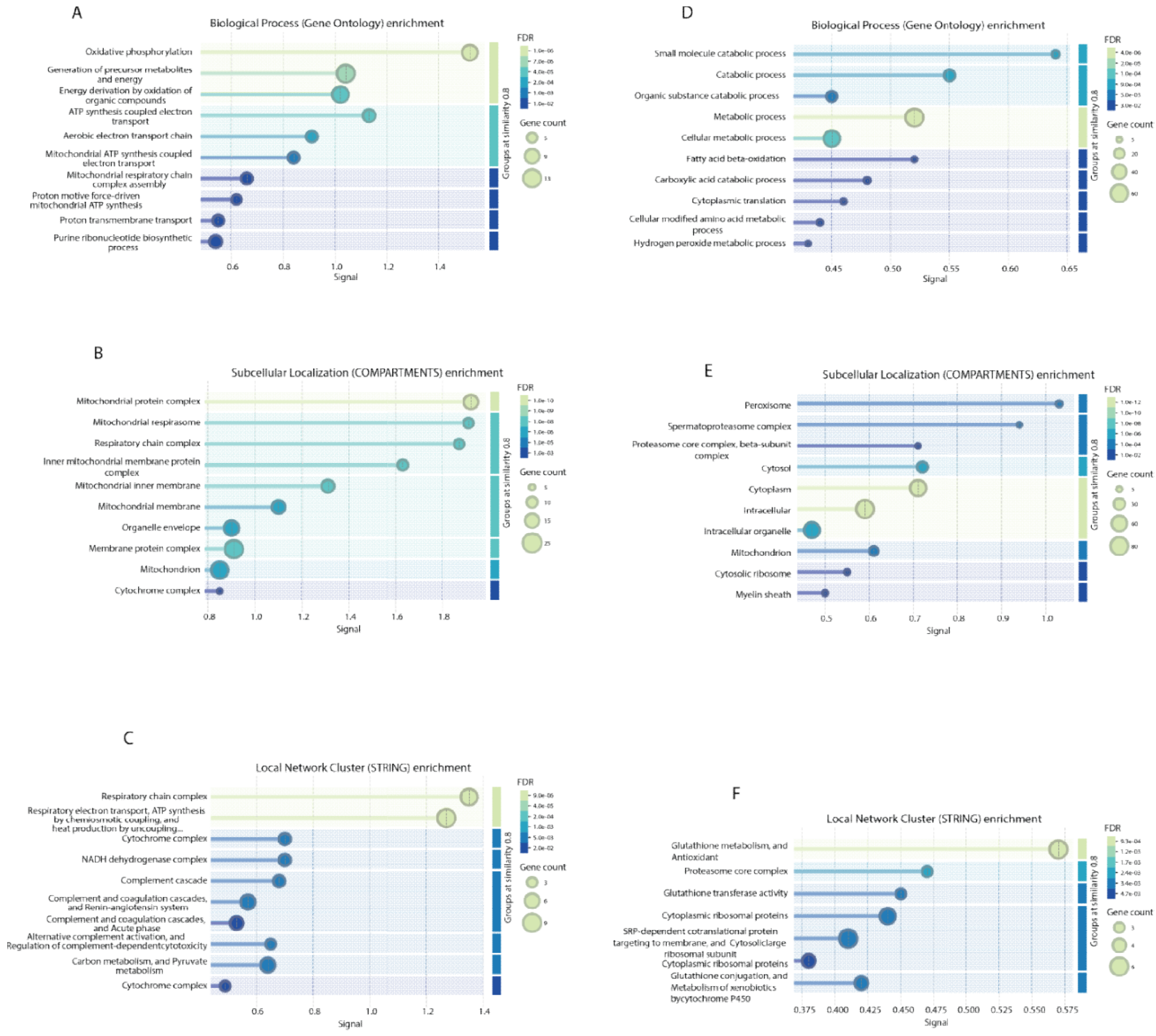


**Figure Supplementary 2. Functional enrichment and subcellular localization of cluster-specific proteins:** Bubble plots showing Gene Ontology (GO) biological process enrichment, subcellular localization (COMPARTMENTS), and STRING network cluster enrichment for proteins derived from cluster 1 (left panels; upregulated/restored by DVA) and cluster 2 (right panels; downregulated by DVA). The top panels represent enriched biological processes, highlighting mitochondrial energy metabolism and oxidative phosphorylation in cluster 1, and catabolic and metabolic processes in cluster 2. Middle panels show subcellular localization, indicating strong mitochondrial enrichment for cluster 1 proteins and broader cytoplasmic, proteasomal, and vesicular localization for cluster 2 proteins. Bottom panels depict STRING-based network clusters, revealing tightly interconnected functional modules corresponding to respiratory chain complexes in cluster 1 and metabolic and proteostasis-related networks in cluster 2. Bubble size reflects gene count and color indicates statistical significance (FDR).
